# Searching match for single-cell open-chromatin profiles in large pools of single-cell transcriptomes and epigenomes for reference supported analysis

**DOI:** 10.1101/2021.03.24.436738

**Authors:** Shreya Mishra, Neetesh Pandey, Smriti Chawla, Debarka SenGupta, Kedar Nath Natrajan, Vibhor Kumar

## Abstract

The true benefits of large datasets of the single-cell transcriptome and epigenome profiles can be availed only with their inclusion and search for annotating individual cells. Matching a single cell epigenome profile to a large pool of reference cells remains a major challenge. We developed a method (scEpiSearch) to resolve the challenges of searching and comparing single-cell open-chromatin profiles against large pools of single-cell expression and open chromatin datasets. scEpiSearch is more accurate than other methods when comparing single cell open-chromatin profiles to single-cell transcriptomes and epigenomes. scEpiSearch also provides a robust method for reference-supported co-embedding of single-cell open chromatin profiles. In performance benchmarks, scEpiSearch outperformed multiple methods for the low dimensional co-embedding of single-cell open-chromatin profiles irrespective of platforms and species. scEpiSearch works with both reference single-cell expression and epigenome profiles, enabling classification of single-cell open-chromatin profiles. Here we demonstrate the unconventional utilities of scEpiSearch by applying it on single-cell epigenome profiles of K562 cells and samples from patients with acute leukaemia to reveal different aspects of their heterogeneity, multipotent behaviour and de-differentiated states. Applying scEpiSearch on our single-cell open-chromatin profiles from embryonic stem cells(ESCs), we identified ESC subpopulations with more activity and poising for endoplasmic reticulum stress and unfolded protein response. Thus, scEpiSearch solves the non-trivial problem of amalgamating information from a large pool of single-cells to identify and study the regulatory states of cells using their single-cell epigenomes.The true benefits of large datasets of the single-cell transcriptome and epigenome profiles can be availed only with their inclusion and search for annotating individual cells.

## Introduction

Single-cell epigenome profiling enables the identification of active and poised cis-regulatory sites and the underlying genome regulation across *in vivo* and *in vitro* cell types, tissues. Due to several advantages like lack of RNA degradation digital read out of chromatin status and better understanding of heterogeneity in cellular responses, single-cell epigenome profiling is increasingly adapted for atlas-scale datasets and provides accurate insights into underlying cell state regulation (1, 2). Hence, an important challenge is how to handle the challenge of searching, and meta-analysis of single-cell epigenome profiles. A search tool can handle such tasks, which can also reveal various stages of de-differentiation of cancer cells and predict a cell’s behaviour in an unknown state. Such an approach can lead to better annotation and regulatory-inference from new single-cell open-chromatin profiles as it leverages the additional information present in reference pool of cells in different cellular-states. The challenges and opportunities of such approaches have been highlighted as one of the eleven grand challenges in single-cell data science by Lahnemann et al.(3), under the topic of mapping a single-cell to reference atlas. They have also listed integrating single-cell data across samples and experiments as another grand challenge. There have been efforts from several (4) groups(5, 6) to build search-engine for single-cell expression profiles. However, they do not resolve the issues associated with such challenges for single-cell epigenome datasets. Whereas, tools like scfind(4) help to search single-cell profiles to identify both cell-type specific and housekeeping genes. There have been a few studies on integrating single-cell epigenome with single-cell expression profiles(5–8), but they have not used the approach of searching a large pool of reference cells. Such tools also include Seurat(9), LIGER(10) and Conos(11), which have been proposed for the integration of single-cell open-chromatin profiles. However, a recent benchmarking study by Leucken et al.(12), compared more than 30 such single-cell integration methods and revealed that most integrative approaches performed poorly for batch correction while integrating scATAC-seq profiles. Leucken et al. revealed a fact about such integrative methods that only 27% of their integration outputs for scATAC-seq profiles performed better than the best-unintegrated results (12). Therefore, despite the availability of large single-cell epigenome atlases(7–9) and methods for their low-dimensional visualization, there is a scarcity of a robust mapping method which can lead to development of search engines meant to correctly match query single-cell epigenome profile to large number of single-cell profiles irrespective of batch effect.

Most of the published approaches that utilize canonical correlation and principal component analysis focus on visualization and analysis of single-cell ATAC-seq profiles within a group, and can miss rare cells (very few cells across the entire dataset). On the other hand, a search engine approach can help in handling every single-cell independent of each other would provide the tremendous benefit of preserving the information of rare and unique cells in a study as well as utilizing the datasets from other studies. In comparison to the single-cell transcriptome, single-cell open-chromatin profiles offer new obstacles. Currently, single-cell epigenome profiling mainly aims to capture open chromatin regions with (preferentially at promoters, enhancers etc.,) using DNase-seq (DNase I hypersensitive site sequencing)(13), MNase-seq (Micrococcal Nuclease digestion with deep sequencing)(14) or ATAC-seq (Assay for Transposase-Accessible Chromatin using sequencing)(15) The compiled read-count matrices using single-cell open-chromatin have more genomic loci (peaks) as features than a similar matrix for single-cell expression dataset. Most often, genomic loci (peaks) in the read-count matrices of single-cell open chromatin profiles compiled by different research groups are not the same. Hence existing algorithms and search methods proposed for single-cell expression profiles cannot be used directly for single-cell open-chromatin profiles.

Here we describe scEpiSearch, consisting of novel computational methods to match queried single-cell open-chromatin profiles with a large pool of reference single-cell open-chromatin and single-cell expression datasets. scEpiSearch resolves the issue of handling non-similar peak-lists of single-cell epigenome profiles from multiple scientific groups and solves the problem of calculating the statistical significance of the match of the query with single-cell expression and open-chromatin profiles. scEpiSearch uses a gene-enrichment score as a proxy for cell-type specificity (instead of gene-activity (15)) while minimizing the bias and noise bias across reference cells (see Methods). scEpiSearch also resolves the problem of using reference cell atlases for highly efficient joint-embedding of query single-cell open-chromatin profiles irrespective of batch effect, species and peak-list. Here we apply scEpiSearch to single-K562 and embryonic stem cell epigenomes and capture heterogeneity, lineage bias and stress-response across single-cells to have a better understanding of regulatory behaviours through their epigenomes.

## Results

scEpiSearch first preprocesses the reference pool of single-cell epigenome and expression profiles (see Methods). In fact, it also has its own processed reference pool of single-cell transcriptomes and epigenomes (Figure 1). Current reference pool of scEpiSearch has 3.3 million expression profiles from human and mouse cells (supplementary Table-1), and approximately 8,00,000 cells from human and mouse cells (supplementary Figure S1). To handle such a large reference pool of single-cell expression and epigenome profiles scEpiSearch keeps it in a clustered format so that they could be searched in a hierarchical manner (supplementary Figures S2, Figure 1).

**Figure 1.**
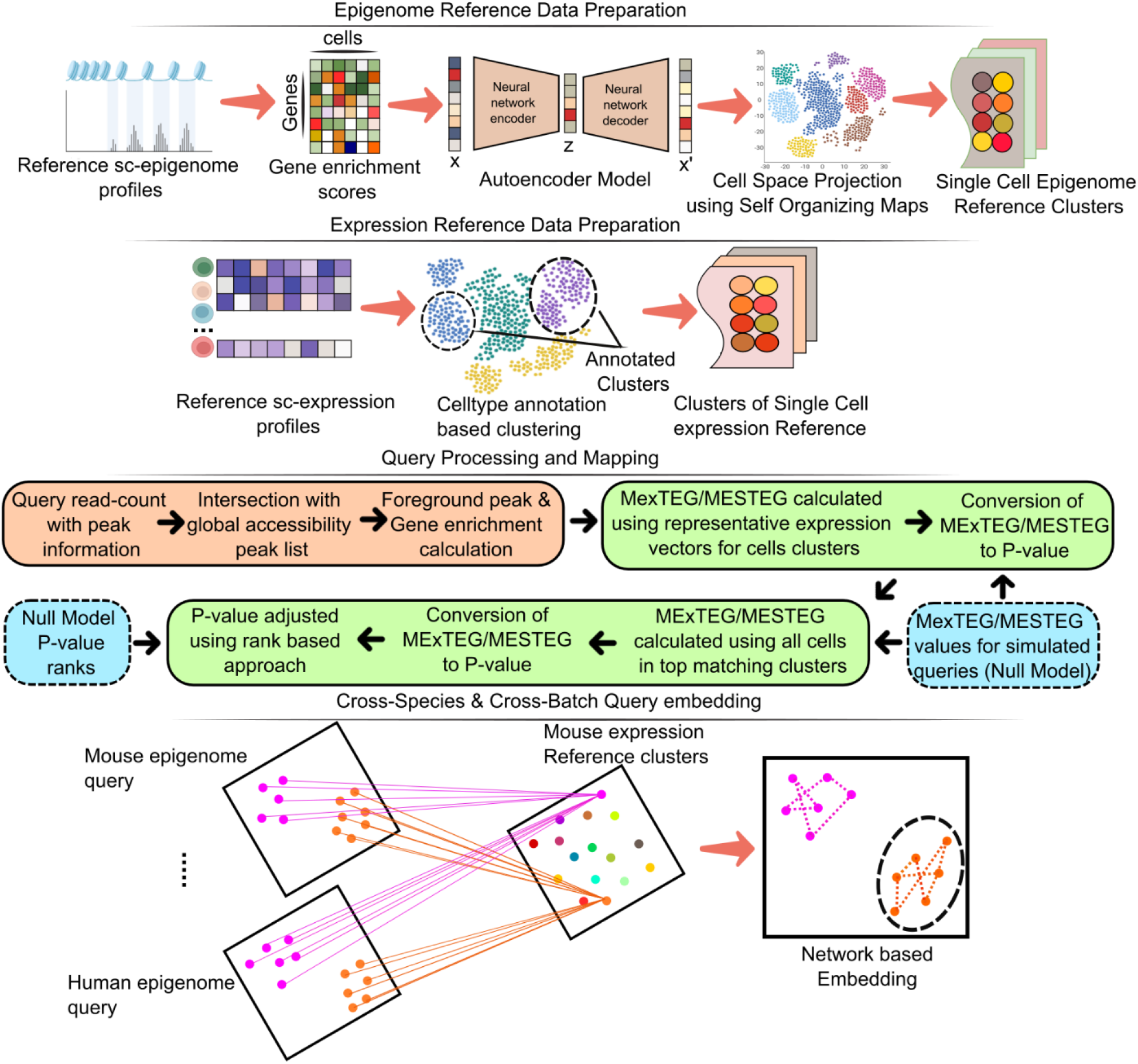
A graphical outline describing the proposed approach and algorithms in scEpiSearch for annotation of new single-cell open-chromatin profiles and better inference of their regulatory state using the collection of available data-sets of single-cell epigenomes and transcriptomes. It involves the steps named as : Epigenome and Expression Reference data preparation, Query processing and Mapping and Cross-species and Cross-batch Query embedding. The cross-species and cross-batch query embedding represents co-embedding of multiple open-chromatin profiles irrespective of difference in peak-list in read-count matrix, batch effect and species using existing reference single-cell profiles.

For both query and reference single-cell open-chromatin profiles, scEpiSearch first normalizes the read-count of every peak with a global accessibility score to highlight potential enhancers(16). For both species, human and mouse, we used the global accessibility peak-list compiled using several published open-chromatin profiles of bulk samples published by different groups and consortiums (17, 18). Normalization by global accessibility score for peaks removes the bias, which could have come from others cells in the same query. Thus, every cell in the query is treated independently of each other. In the faster version of scEpiSearch, predefined peak list is used to determine the proximal genes directly. For a query with more than 200000 peaks, intersection with a predefined peak-list with global accessibility speeds up the process of determining proximal genes by 1000 times (Supplementary Figure S1c) with almost 75-90% peaks covered most of the time. (Supplementary Figure S1c-d).

### scEpisearch enables correct matching to reference cells irrespective of technical biases

scEpiSearch uses peaks with high normalized counts as foreground and others as the background set to calculate the gene-enrichment score using the Fisher exact test (hypergeometric test). For matching to a reference single-cell transcriptome profile, it uses its normalised expression (see Methods) values to calculate median expression for the top 1000 enriched genes (MExTEG) of query cells. For a query, the MExTEG of a reference cell is converted to P-value using precalculated MExTEG values for cells in the null model (see Figure 1 and Supplementary Figure S2). In order to compare to a large pool of reference single-cell transcriptome profiles scEpiSearch uses a hierarchical approach, it first matches the query with representative expression vectors for clusters of cells. Then reference cells from top matching clusters are used to calculate MExTEG values and corresponding P-values. scEpiSearch further calculates a new P-value based on the ranks of reference cells for a query to reduce bias in the dataset and search procedures (see Methods, Supplementary Figure S2). In other words, scEpiSearch makes a rank adjustment for hits using their precalculated ranks for the null model. First, we compared our method using a reference set of 10000 mouse single-cell expression profiles from mouse cell atlas (MCA). We compared it against 3 different approaches: i) comparing gene-activity of scATAC-seq to gene-expression, ii) correlating gene-enrichment scores of scATAC-seq and expression profile of reference iii) calculating correlation between BABEL based predicted expression of query scATAC-seq profile to reference expression (Figure 2A). We found that our MExTEG based approach is much superior to direct comparison (or correlation) of reference gene-expression values to gene-activity, gene-enrichment or predicted expression of query scATAC-seq profiles (Figure 2A).

**Figure 2:**
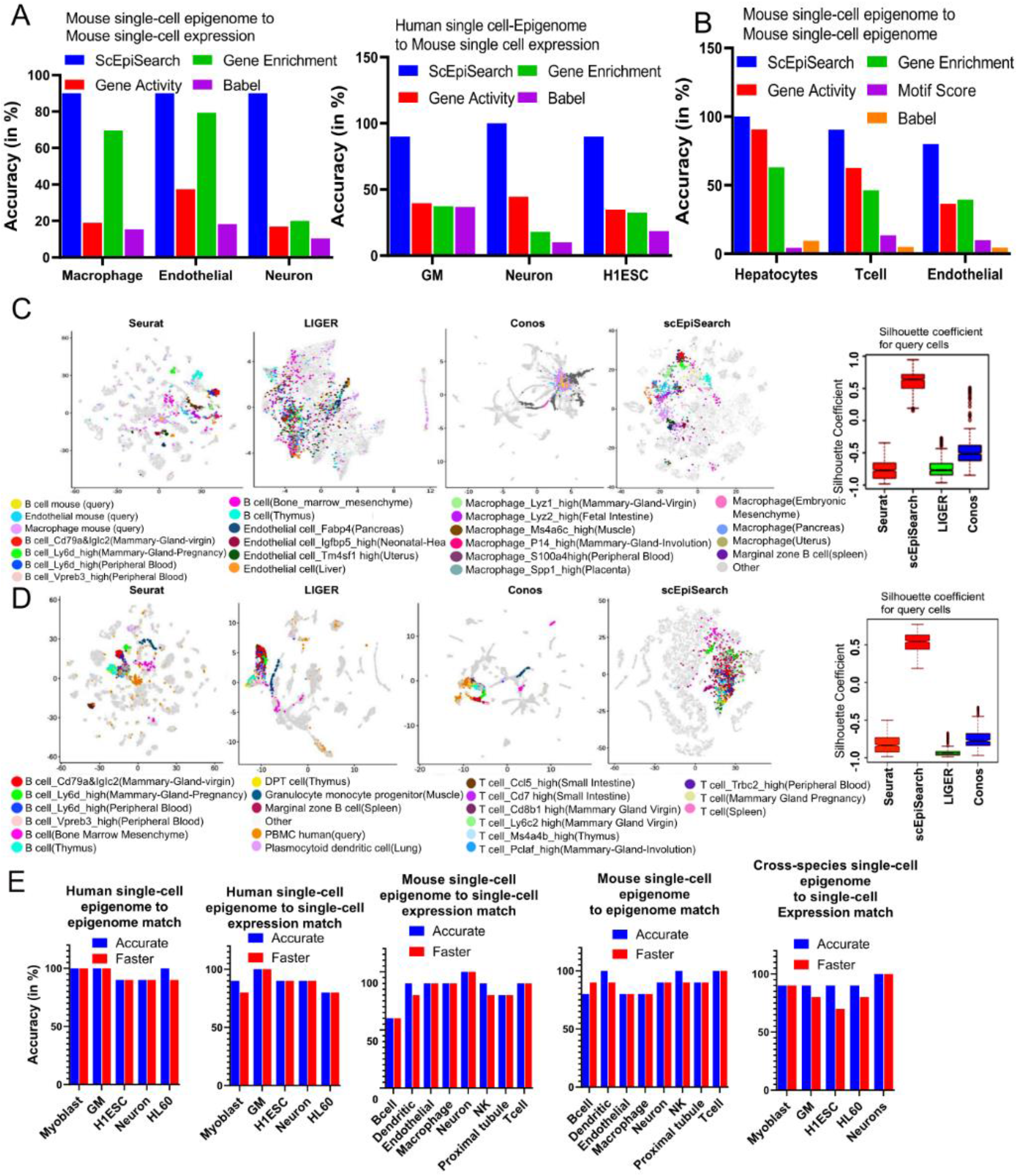
Evaluation of the accuracy of scEpiSearch. **A)** Comparison of scEpisearch with 3 approaches based on correlation of extracted gene scores for matching single cell open-chromatin profiles to a pool of reference single-cell transcriptome. Here the reference dataset consisted of single-cell expression profiles of 10000 cells chosen from mouse cell atlas (MCA). Accuracy here shows the percentage of query cells which had correct cell-type among top 5 top matches. **B)** Comparison of 5 methods for matching query single cell open-chromatin profile to reference sci-ATAC-seq profile of cells chosen from data-set published by Cusanovich et al. **C)** Comparison of scEpiSearch with integrative method using reference single-cell expression profiles of 10000 cells from mouse cell atlas (MCA) dataset. Here, query consisted of scATAC-seq profiles of 3 types of mouse cells namely; B cell, macrophages and Endothelial. The silhuotte index of query cells for being in proximity to correct reference cell-types, is shown for different methods. **D)** Evaluation of cross-species search for integrative methods and scEpiSearch using human PBMC scATAC-seq profiles as query and reference single-cell expression profiles from MCA. Silhouette coefficients for human PBMC cells are shown for different methods. Here immune cells in references and query cells were considered to belong to one class, where as other cell-types as second class for calculation of silhouette coefficients. **E)** Accuracy achieved with scEpiSearch engine for query scATAC-seq read-count matrices shown from left to right as such: i) query human single-cell epigenome (open-chromatin) profile to reference Human single-cell epigenome ii) query human single-cell open-chromatin profile to reference human single-cell expression collection iii) query mouse single-cell epigenome to reference single-cell expression profiles. iv) query mouse single-cell epigenome (open-chromatin) to reference mouse single-cell expression profile v) cross-species search – query human single-cell epigenome to reference mouse single-cell expression. Y-axis shows accuracy in the percentage of query cells for which correct annotation came among the top 5 hits. It shows accuracies as bar-plots for faster and accurate modules of scEpiSearch.

For matching to a reference scATAC-seq profile, its median enrichment score for top enriched genes (MESTEG) of query cells is calculated. The MESTEG value is converted to P-value using precalculated MESTEG scores for cells (vectors of enriched genes) in the null model (Methods, Figure 1, Supplementary Figure S2). ScEpiSearch also uses a hierarchical approach to find the most matching scATAC-seq profile in the reference dataset (Supplementary Figure S2). After determining the rank of reference cells for a query cell, it calculates a new P-value using the precalculated rank of the same reference for the null model (see Methods). We compared MESTEG based approach to direct correlation between gene-activity, gene-enrichment and BABEL based predicted expression of query and reference open-chromatin profiles. In our evaluation query scATAC-seq profiles were not from the same data-sets which was used make reference pool of single-cell epigenomes. MESTEG based approach was substantially better than other compared procedures in term of finding correct matching open-chromatin profiles.

We also compared scEpiSearch to integrative methods, Seruat(9), LIGER(10) and Conos(11) in terms of finding the closest matching single-cell expression profile for various type of scATAC-seq read-count matrices. Here we used the same reference single-cell expression pool of ∼10000 cells from the MCA dataset (19). We provided the same reference cells to Seruat(9), LIGER(10), Conos(11) and scEpiSearch. For 2D visualization of scEpiSearch results, we used the average of coordinates (in tSNE plot) of top 3 matching cells. We found that with homogeneous and smaller query set of single-cell open-chromatin profiles, the co-embedding results by integrative methods were not satisfactory (see Supplementary Figure S3). It could be due to the design of integrative methods like Seurat and LIGER to exploit heterogeneity in single-cell profiles to find anchors, resulting in the wrong grouping of homogenous query single-cell ATAC-seq profiles with various non-similar cells. In fact, Leucken et al. (12) also revealed poor performance of integrative methods for scATAC-seq data and mentioned that gene activity scores used by many integrative methods could be poorly suited to represent scATAC-seq data. One another reason for the revelation of such a result could be that, unlike previous studies, we did not calculate silhouette coefficients/index for reference single-cell expression data-points in the co-embedding plots, as it could have overwhelmed the corresponding values for query single-cell ATAC-seq profiles (as shown in supplementary Figure S3c). Our target was to evaluate the process of finding matching expression profiles for single-cell ATAC-seq data-sets (like a search engine); hence we calculated silhouette coefficients only for query cells.

The co-embedding plot improved when we increased heterogeneity in the query to include the single-cell ATAC-seq profiles of 3 types of mouse cells (macrophages, B cells and endothelial cells)(Figure 2C). However, based on the measure of silhouette coefficients for query single-cell ATAC-seq profiles, the integrative methods (Seurat, LIGER and Conos) were not comparable to scEpiSearch (Figure 2C). A similar trend was observed when scATAC-seq profiles of human cells were used as a query for the same reference expression dataset consisting of 10000 cells from MCA. When scATAC-seq profiles of Human embryonic stem cells (HESC) were used as a query, scEpiSearch based plots showed them closer to mouse ESC. Similarly, for the scATAC-seq profile of Human Neuron cells, results based on scEpiSearch showed their proximity to reference neuronal cells from MCA (supplementary Figure S4). Whereas Seurat and LIGER had the same problem, such that homogenous query cells were spread and colocalized with multiple groups of non-similar reference expression profiles in co-embedding plots (see Supplementary Figure S4). When we used a query consisting of human PBMC scATAC-seq profiles with higher heterogeneity, the results for LIGER and Seurat improved such that query cells appeared closer to immune cells in their co-embedding plots (Figure 2D). However, when we calculated the silhouette coefficients for query cells to evaluate the efficacy of the integrative method as a search engine, their performance was not comparable to simple embedding based on the results of scEpiSearch. Overall, our results indicate that integrative methods could not provide an efficient search for matched expression profiles for query scATAC-seq datasets like scEpiSearch.

The standalone and webserver versions have their own database of reference expression and gene-enrichment scores of scATAC-seq profiles. Both versions of scEpiSearch have scalable visualization, where results of more than 1000 query scATAC-seq profiles can be visualized interactively. Evaluation using several scATAC-seq profiles with known cell-type annotation revealed that for most of the queries, the accuracy of the search of scEpiSearch is about 80-100% in highlighting the correct cell type among the top 5 matches (Figure 2E, supplementary Tables 2-3). Our approach of using proximal genes enrichment and expression profiles, rather than relying on correlation or distances among cells, also helps in avoiding artefacts due to technique or species-specific batch effect. Therefore, scEpiSearch also allows matching query scATAC-seq profiles from human cells to mouse reference single-cell expression profiles with high accuracy (see Figure 2E, supplementary Table-4).

There are several possible applications of scEpiSearch, including i) finding cell types for scATAC-seq profiles from unannotated cells from *in vivo* samples ii) Studying heterogeneity and tracking divergences in state of cells and their potency, for example, one cell-line showing differentiation potential towards several lineages, such as K562 cells(20) iii) Finding matching mouse model for a human cell which is not characterized well. iv) Highlighting marker genes representing query cells v) Embedding and clustering multiple scATAC-seq profiles irrespective of their sources, technique and species. Here we have applied ScEpiSearch for four types of case studies to demonstrate such applications.

Further, applying scEpiSearch to find the closest match for “unknown” cells in the single-cell indexed ATAC-seq (sciATAC-seq) dataset published by Cusanovich et al.. provided very relevant hits. Such as for the sciATAC-seq profile of unknown cells from mouse liver cells, 96% of the top 5 matching scRNA-seq profiles were from hepatocytes (Supplementary Figure S5) and majority of unknown cells from prefrontal cortex had top matching cell-type as neurons. scEpiSearch also provides the list of gene enrichment scores for query cells to help researchers decide on representative genes (or markers) or develop the reliability of results. Hepatocytes markers such as like Apob, Tat, Cfhr2, Mat1a were present frequently among the top 20 enriched genes for “unknown” query cells from the liver (Supplementary Figure S5a). We have provided some more examples in supplementary Figures S6 and S7. Application of scEpiSearch for finding matched reference single-cell expression profiles for scATAC-seq read-count matrixes of peripheral blood mononuclear cells (PBMC) from healthy humans, provided top matches in concordance with the expected distribution of cell types in the blood (supplementary Figure S8a) (21).

### Determining lineage of cancer cells and understanding their multipotent behaviour using scEpiSearch

Recently several groups have started profiling scATAC-seq profiles of frozen nuclei derived from tumour samples. Tumour cells are known to have heterogeneity and often appear to have unreported intermediate cellular states. Reprogramming and de-differentiation in cancer cells are often associated with drug resistance and unexpected lineage switching(22, 23). Thus, it becomes important to compare the scATAC-seq profile of cancer cells to the existing pool of cells to find their lineage and potency for a better understanding of tumour pathogenesis. Hence as proof of concept, we first evaluated the performance of scEpiSearch in identifying lineage using scATAC-seq profiles of HL60 and K562 cell lines. HL-60 cells derived from myeloid leukemia patients show neutrophilic promyelocytic morphology(24). For scATAC-seq read-count matrices of HL60 cells, scEpiSearch found that top matching expression profiles were from myeloid lineage cells (dendritic and Langerhans cells). For 10% of the HL60 cells, scEpiSearch also reported monocytes among the top 3 matching cells (Figure 3A, supplementary Table-5). Such results could be due to differentiation potential of HL60 towards monocyte which is well known(25). The most frequently enriched genes included LYST, ALOX5, RASSF4, AOAH, RXRA, MEF2A, PRAM1 and AKAP13 (Supplementary Figure S9a), which have been catalogued in gene-sets for myeloid lineage in Human_Gene_Atlas listed in EnrichR(26) (https://maayanlab.cloud/Enrichr/#stats)

**Figure 3:**
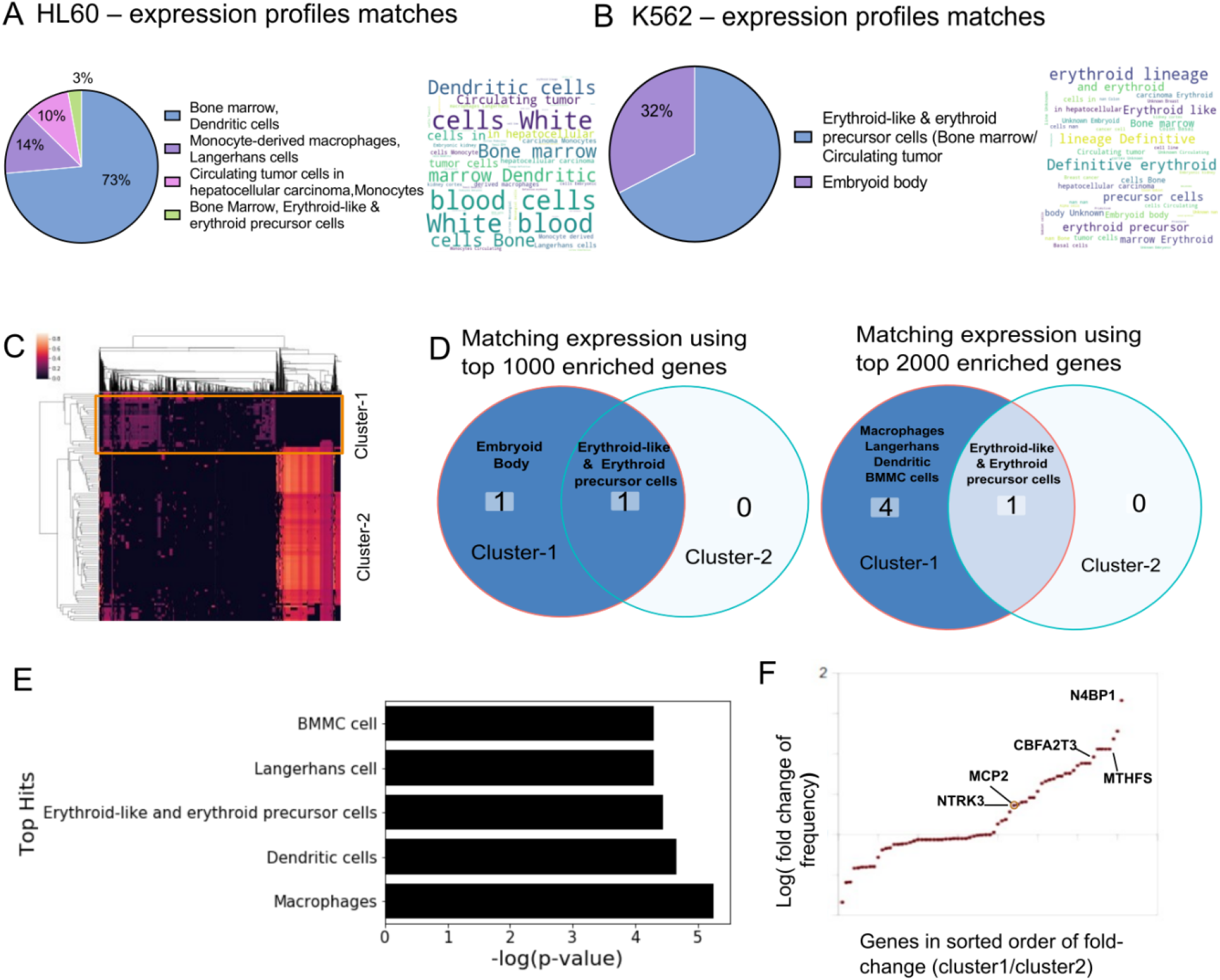
Case study of using scEpiSearch to reveal lineage and underlying multipotency of cancer cells. **(A)** The pie chart shows the proportion of cell types for top 3 matching single-cell expression profile for scATAC-seq read-count matrix for HL60 cells. A word-cloud of annotations of matching single-cell expression profiles is also shown on the right-side of the pie-chart. B) The proportions of cell types for top 3 matching single-cell expression profiles for scATAC-seq read-count matrix for K562 cells. The corresponding word-cloud is shown on the right. (C) Heatmap of epigenome matching profiles is shown. X-axis shows matching epigenome profiles from reference, and Y-axis shows Query cells. Two clusters of query cells are also shown. D) The annotations of cells for the top 3 match expression profiles matching to scATAC-seq dataset of K562 cells in cluster-1 and cluster-2. The result has been shown when top-1000 and top-2000 enriched genes are used. E) Average P-values for matching to cluster-1 K562 cells with different types of cells. **F)** Fold change in frequency of being in top 50 enriched genes is shown for query K562 cells from two clusters. Each dot represents a gene. It is plotted only for those genes which have a frequency of at least 10% in either class and have a fold change above 1.05. Genes whose names are displayed in the plot are known to be markers for dendritic lineage.

The K562 leukaemia cell line has been broadly utilized in consideration of erythroid differentiation. K562 cells serve as an experimental model to study the early steps of megakaryoblast and macrophage commitment and differentiation (20, 27). Using default parameters of using top 1000 enriched genes in scEpiSearch, the top 3 hits for all K562 cells consisted of Erythroid-like and erythroid precursor cells (Figure 3B). In top-3 matching expression profiles, we also found a few cells from the embryoid body whose lineage was not annotated. Genes with a higher frequency of being among the top 50 enriched genes for query scATAC-seq profiles of K562 included CES5A, DAOA, KSR1 and PRKCB (supplementary Figure S9b), which have been reported to have higher expression in erythroid cells in the mouse cell atlas scRNA-seq dataset by Han et al. (19). At the same time, frequent top enrichment of genes involved in early erythropoiesis like NR2F2 and SOCS1 (28, 29) hints at the de-differentiated state of K562 and their similarity with erythrocyte progenitors. Overall, scEpiSearch is able to predict the major lineage of cancer cells and is useful in highlighting relevant genes.

Open chromatin profiling also captures regions that are poised for future activity during differentiation. Therefore we aimed to study the multipotent behaviour of K562 cells using their scATAC-seq profile with the help of scEpiSearch. For detecting multipotency and heterogeneity among K562 cells, we used the clustering result provided by scEpiSearch, which is based on their match with the reference epigenome profile. Clustering of K562 scATAC-seq profiles revealed two major clusters (Figure 3C). We compared results of matching expression profiles found by scEpiSearch using top 1000 and top 2000 enriched genes for the query. Cells in cluster-2 always had top 3 matching expression profiles from erythroid like or erythroid precursor cells with both parameters settings (top-1000 and top-2000 enriched genes) (Figure 3C-D). However, cells in cluster-1 also had other types of matching expression profiles in addition to erythroid-like cells. Using top-2000 enriched genes of query cells, other matching expression profiles for cluster-1 cells were Macrophages, dendritic cells (including Langerhans) and a few bone marrow mononuclear cells (BMMC) (Figure 3D). We took an average of p-values for top-3 expression matches provided by scEpiSearch for k562 cells from cluster-1. We found that expression profiles of macrophages had the most significant p-values of the match with cluster-1 cells (Figure 3E). For dendritic and erythroid lineage cells, average p-values for the match to cluster-1 cells were also significant but comparable to each other. We also found a few genes labelled as markers of dendritic cells (available in EnrichR (26)) like NTRK3, MCP2, MTFHS, N4BP1, CBFA2T3, which had a higher frequency of appearance in the top 50 genes for cluster-1 cells in comparison to cluster-2 (Figure 3F). The differentiation potential of K562 cells towards macrophage lineage is well known(27). Several groups have also reported the potency of K562 cells to differentiate toward dendritic lineage(30). Our analysis revealed that a minor population of K562 cells had slightly more significant similarity with macrophages than erythroid lineage. It also hints toward a potential application of scEpiSearch in studying heterogeneity in the underlying poised state of cancer cells for oncology studies.

### scEpiSearch enables joint embedding and visualization of single-cell epigenome profiles across batches and species

Even though scEpiSearch finds matching transcriptome and open-chromatin profiles for single query cell epigenome, it is often necessary to visualize and cluster cellular profiles from multiple sources together to get an insight into discrete or mixed cellular states. Therefore scEpiSearch is also designed to embed and provide an integrated visualization of multiple scATAC-seq profiles with different peak-list and batch-effect irrespective of their species of origin. scEpiSearch calculates distances among query cells based on the similarity of top-matching mouse reference expression profiles. Here, mouse expression profiles are called similar if they belong to the same cluster in the processed reference scRNA-seq dataset of scEpiSearch. We compared the performance of scEpiSearch with four methods meant for embedding (SCANORAMA, MINT, SCVI, SCALE)(31–34) using four different collections of scATAC-seq read-count matrices. SCANORMA, MINT and SCVI use genes as features hence gene-enrichment scores from multiple scATAC-seq profiles (query cells) were provided to them for 2D embedding. Whereas for SCALE, its latent space representation of scATAC-seq read-count matrices was used with tSNE to perform 2D embedding. As shown in Figure 4 and Supplementary Figure S10a, the 2D embedding plot made by scEpiSearch for scATAC-seq profiles has almost correct colocalization of similar cell types irrespective of the species and laboratory of origin. Other available methods of embedding (SCANORAMA, MINT, SCVI and SCALE) provided the wrong grouping of cells (Figure 4 and Supplementary Figures S10a). For further confirmation, we estimated clustering purity after density-based spatial clustering (using DBSCAN(35)) of the embedding results using cell type labels as true clusters. The clustering purity based ARI (adjusted Rand Index) and NMI (normalized mutual information) scores showed the superiority of scEpiSearch in the embedding of open-chromatin profiles in an unbiased manner (Figure 4, Supplementary Figure S10a). The silhouette coefficients -based comparison is shown in Supplementary Figure S10b.

**Figure 4:**
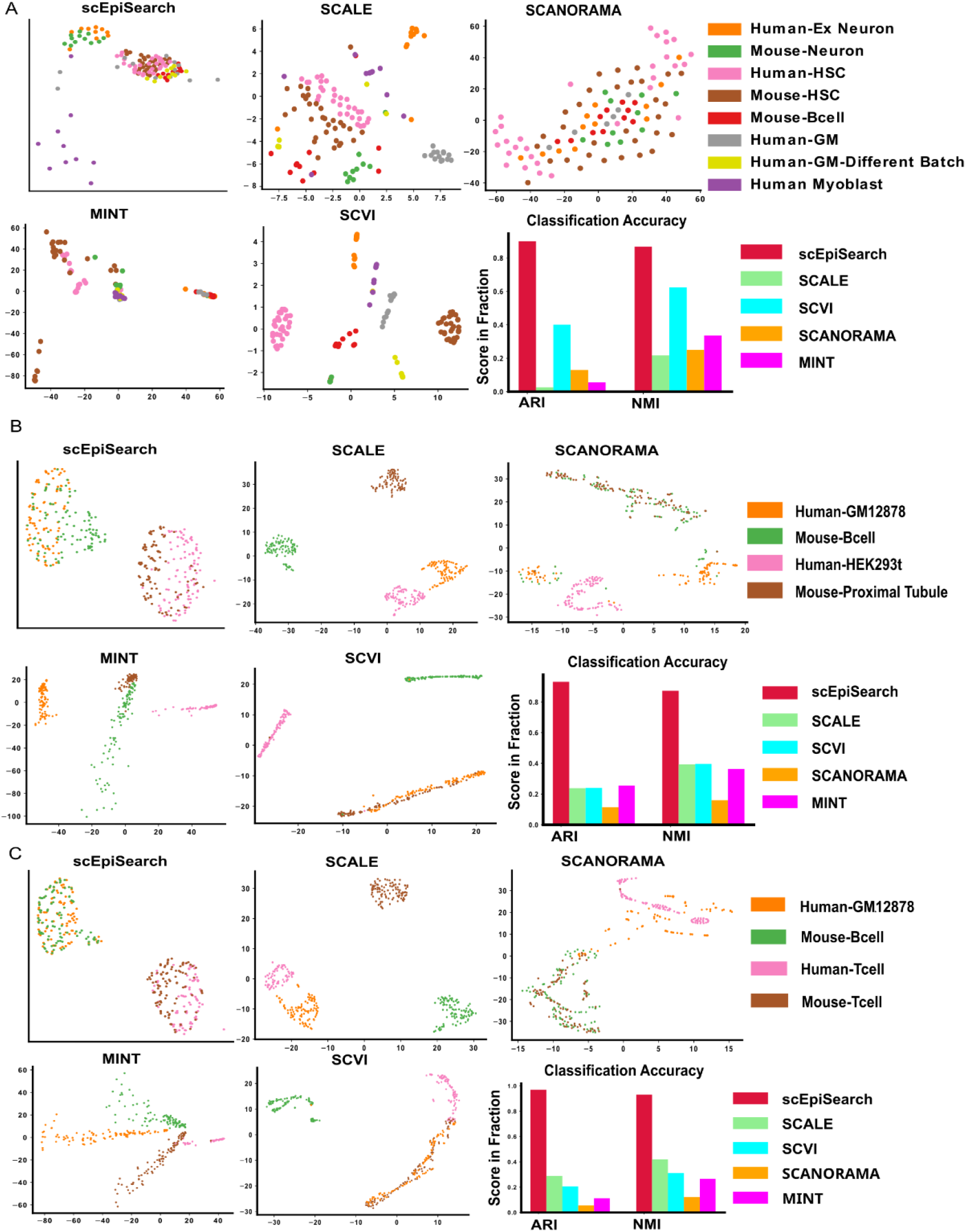
Evaluation of embedding of query sets of single-cell open-chromatin profiles irrespective of batch effect, species, differences in peak-list and their source (scientific group). (A) For this case study, queries consisted of separate read-count matrices for scATAC-seq profiles of human-neuron, mouse-neuron, human-HSC (hematopoietic stem cells), house-HSC, human-Myoblast, human-GM12878 (GM) cells from two batches and mouse-B-cells. The peak lists of read-count matrices were different from each other. Embedding plots from 5 different methods are shown here. While SCANORAMA mixed the location of all cells, MINT could not group cells of the same type together like scEpiSearch. The purity of density based spatial clustering (using DBSCAN) with embedded coordinates is also shown here in terms of ARI (Adjusted Rand Index) and NMI (normalized mutual information) scores. The silhouette coefficients calculated without DBSCAN based clustering are shown in Supplementary Figure S10b. (B) The plot of embedding results shows the alignment of the same cells from different species and batches together. Queries are made for Human-GM12878 cell, Mouse B-cell, Human-HEK293T, Mouse-Proximal tubule. Embedding plot from ScEpiSearch derived from projections onto mouse expression profiles. (C) The plots show 2D embedding of cells from different species and batches. Queries were made for Human-GM12878 (GM), Mouse B-cell, Human T-cell, Mouse T-cell.

### A case study of embedding: understanding multiple phenotype acute leukaemia

To get further insights from the joint embedding of single-cell epigenome profiles and underlying cell states, we analyzed scATAC-seq profiles from patient blood cells with mixed-phenotype acute leukaemia (MPAL) (36). An initial analysis of MPAL cells and PBMC from healthy patients in the same study revealed a change in the fraction of cell types. For the scATAC-seq profile of PBMC from healthy individuals, the matching expression profiles had similar fractions for different cell types as reported by others (21) (Supplementary FigureS8a). However, for MPAL cells from two patients, we found an increase in the fraction of cell types of dendritic, monocyte and erythrocyte lineage (Supplementary Figure S11). It is not trivial to find whether it represented true fractions or it was due to sampling bias during scATAC-seq profiling. Nevertheless, we performed embedding of scATAC-seq profiles of MPAL cells from two patients, PBMC from healthy individuals and progenitors of cells in the blood (progenitors of hematopoietic cells)(37), T cells and B cells (38)(supplementary Methods). For our analysis, we included scATAC-seq profile of progenitors of hematopoietic cells(37) in the blood, namely MEP (megakaryocytic-erythroid progenitor), CMP (common myeloid progenitor), CLP (common lymphoid progenitor), GMP (granulocyte-monocyte progenitor), MCP (mast cell progenitor). In the 2D embedding results from scEpiSearch many MPAL cells overlapped with different types of hematopoietic progenitor cells. PBMCs are grouped together with T cells, B cells and a few MPAL cells (Figure 5A). In our embedding results, PBMC rarely had any overlap with hematopoietic progenitor cells. Such results hint at the different levels and types of de-differentiated states of MPAL cells. The de-differentiated states of MPAL cells could explain their plasticity and lineage switching capability(23). Further detailed analysis revealed that only a few MPAL cells for two patients showed overlap (Figure 5B). The majority of cells from two patients with MPAL did not overlap with each other and showed closeness with different types of hematopoietic progenitor cells. We also applied SCANORAMA, MINT, SCVI, SCALE on the same set of read-count matrices (supplementary methods) and found that they either mixed the location of different types of blood cell progenitors or showed no colocalization of PBMC with B cell or T cell (Figure 5C). Given the fact that B cell and T cell are frequently present among PBMC, it became quite evident that using other methods could not lead to the result achieved by scEpiSearch (Figure 5). In summary, scEpiSearch can display closeness and differences among subpopulations of cells irrespective of source and batch effect and highlight de-differentiated states and plasticity of cells derived from patient samples.

**Figure 5:**
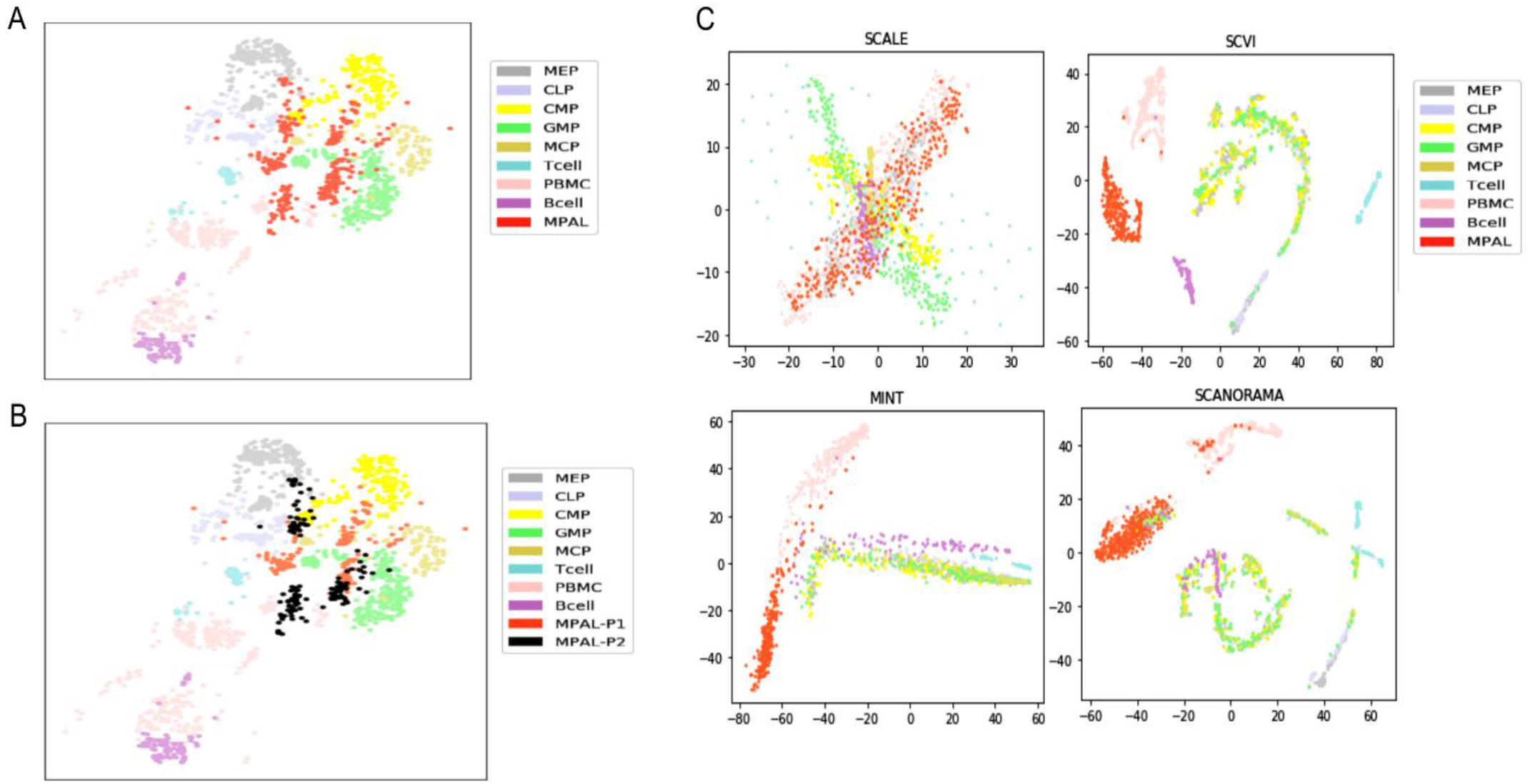
Using scEpiSearch for 2D embedding to track the de-differentiated state of leukaemia cells from blood-cancer patients. **(A)** scEpiSearch based 2D embedding of single-cell open chromatin profiles of 3 types of cells: blood cells collected from patients with mixed phenotype acute leukaemia (MPAL), peripheral blood mononuclear cells (PBMC) from healthy (normal) individuals and progenitors of blood cells. The scATAC-seq profile of MPAL and PBMC cells was published by Granja et al., and progenitor cell epigenome profiles are from a different study. In the embedding plot by scEpiSearch, most of the PBMC cells are far from progenitor cells and closer to B cells. MPAL cells are closer to progenitor cells. Some MPAL cells even completely overlap with blood cell progenitors, highlighting their highly de-differentiated state. **(B)** The same 2D embedding result as above, but now the MPAL cells are coloured according to their patient origins (P1 and P2). Even though MPAL cells from two patients show some overlap, they also have different directions of de-differentiated states. **(C)** Results from other tools for 2D embedding of single-cell open chromatin profile of 3 types of cells: blood cells collected from patients with mixed phenotype acute leukaemia (MPAL), peripheral blood mononuclear cells (PBMC) from healthy individuals and progenitors of blood cells. Other methods either mixed up the locations of different types of hematopoietic progenitor cells or could not colocalize B and T cells with PBMC.

### Application in highlighting unique regulatory patterns in a subpopulation of stem cells

Heterogeneity within individual stem cells has been widely identified in single-cell genomics studies. We hypothesized that clustering scATAC-seq profiles of embryonic stem cells based on matched scores with reference datasets could highlight cells with differential peak enrichment across features. We generated and re-analyzed plate-based scATAC-seq data from mouse embryonic stem cells (mESC) in Serum conditions (cite PMID: **30559361**), applied scEpiSearch combining scATAC-seq profiles and calculated read-counts per cell for clustering of open-chromatin patterns. The hierarchical clustering of our queried scATAC using match score with reference datasets (using top 2000 enriched genes) captured 4 major clusters of mESC cells. The cells in four clusters had high matching scores with reference profiles belonging to published embryonic stem cells, epiblast cells and different blastocyst stage cells (Figure 6A). The cluster-1 cells matched closely to late-blastocyst cells, while cells in clusters 2 and 4 matched with mid and early blastocyst cells (Figure 6B). Using top 10,000 peaks per cluster with the highest normalized read-counts, we performed Gene Ontology (GO) enrichment using GREAT(39), and found cluster-1 cells were enriched for negative regulation of G2/M phase, apoptosis, cellular response to unfolded protein, H4K5 and H4K8 acetylation and DNA damage terms (Figure 6C, supplementary Table-6). In order to have a systematic overview, we selected a few top-enriched terms from each of the four clusters of cells (supplementary Table-6). Then for the selected terms, we curated enrichment scores (P-values) calculated by GREAT for each cluster (see Figure 6C). We found that terms like positive regulation of intrinsic apoptotic signalling pathway were specifically enriched for cluster-1 cells (Figure 6C).

**Figure 6:**
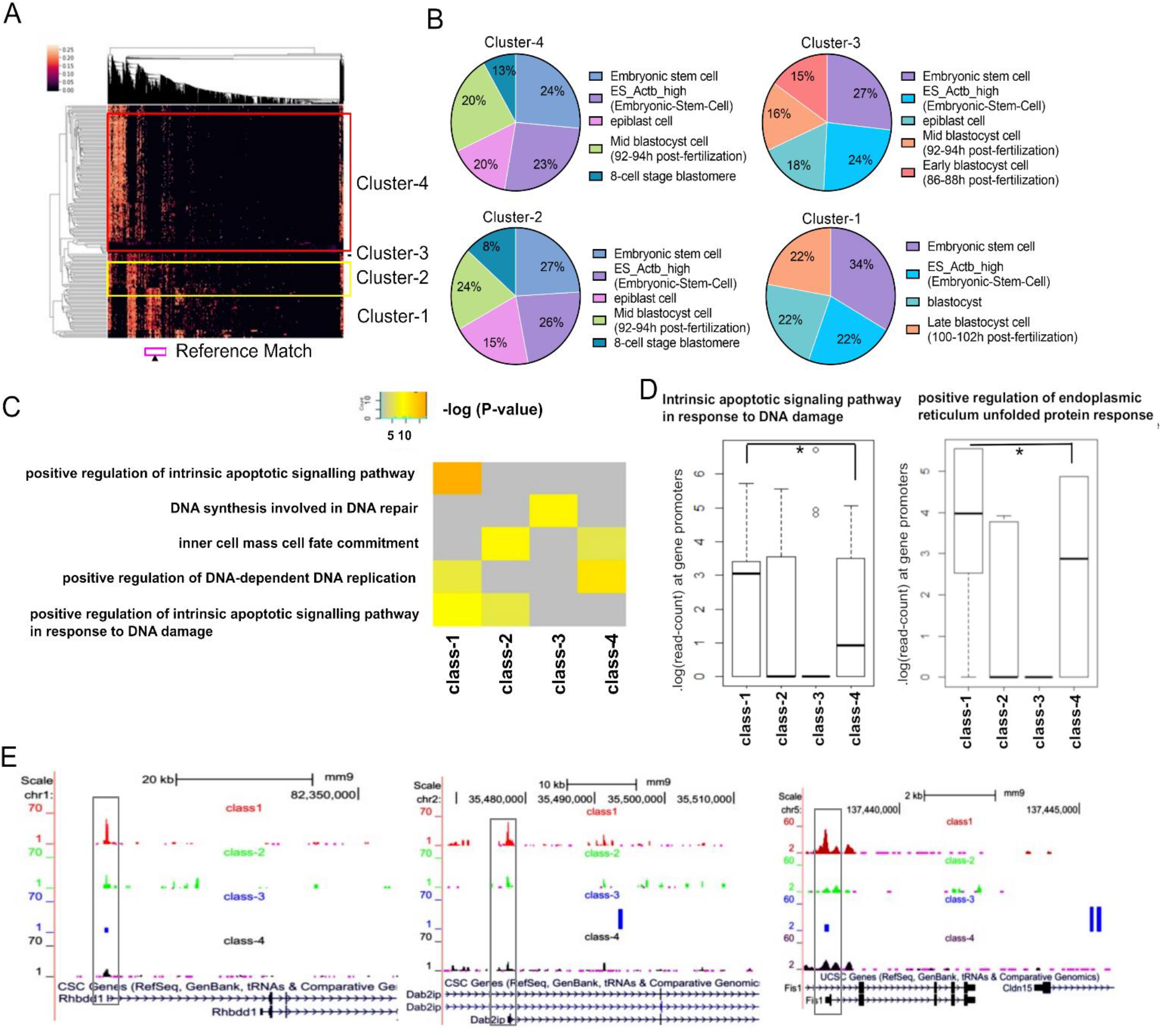
Studying single-cell epigenome profile of stem cells using scEpiSearch. (A) Heatmap obtained from scEpiSearch showing biclustering of match scores of query scATAC-seq profile of mouse embryonic cells (rows) with top matching reference single-cell epigenome profile (columns). The columns, highlighted with the pink box below (and arrow), belong to reference cells from the late blastocyst, which show high similarity with mESC of clusters-1. (B) The pie charts show cell types of top 5 matching expression profiles to query scATAC-seq profiles of mESCs belonging to different clusters. It is based on top-2000 enriched genes based search. (C) The result of gene-set enrichment for selected biological functions using genes proximal to top 10000 peaks specific to mESC cells belongs to different clusters. The gene-set enrichment was performed using the GREAT gene-ontology enrichment tool using default parameters. (D) The read-counts at the promoter of genes belonging to gene-set for biological function terms : “*intrinsic apoptotic signalling pathway in response to DNA damage”* and “positive regulation of *endoplasmic reticulum unfolded protein response”*. Here matrix consisting of read-count for 4 clusters is quantile normalized to avoid any bias. Star(*) shows a significant p-value (< 0.03) calculated using Wilcoxon rank-sum test. (E) The snapshot of UCSC genome browser shows the difference in the activity level of the promoter of two genes (Rhbdd1, Dab2ip) associated with unfolded protein response and endoplasmic reticulum stress and one gene (Fis1) associated with apoptosis, cell cycle and mitochondrion fission.

The gene-set enrichment performed by GREAT(39) often uses genes lying far away from peaks; hence, some more evidence is needed to support its estimation. Therefore, we calculated the read-counts at promoters of all Refseq genes for all four clusters of mESC and performed their quantile normalization. Except for class-3, which had a lower number of cells, we computed normalized read-counts at promoters for other classes and observed that genes associated with *intrinsic_apoptotic_signalling_pathway_in_response_to_DNA_damage*and *ER_unfolded_protein_response* had higher open-chromatin accessibility at their promoters in cluster-1 cells (see Figure 6D). For comparison, we highlight read counts (as box plots) on promoters of other control gene-sets (see Supplementary Figure S12a). We believe that cells in cluster-1 have higher chromatin plasticity poised for cellular responses like apoptosis and ER stress compared to cluster 4 with post-replicated single-cells and defined chromatin state.

We visualized the differences in peak accessibility at single-gene promoters (UCSC genome browser) across clusters. While pluripotency factors (Oct4 and Sox2) had no substantial differences in peak accessibility at promoters (Supplementary Figure S12), we observed higher accessibility in cluster 1 cells for Fis1 related to apoptosis and Rhbdd1 (40)and Dab2ip (41)(Figure 6E) associated with ER stress and unfolded protein response. Our results are consistent with earlier studies describing ER stress due to unfolded proteins in stem cells (42). It also demonstrates a better understanding and interpretation of single-cell open-chromatin peaks can provide novel insights into the heterogeneous stem cell chromatin landscape and underlying regulation.

## Discussion

A major challenge in single-cell genomics is to have an accurate and robust projection of single-cell epigenome profiles to reference epigenome and expression atlases and meaningful interpretation of matching cells. This challenge is confounded by batch and technical biases, including cell-to-cell variability both in signal (peak accessibility) and noise, differential read-depths, protocols, platforms and laboratories. Our proposed computational methods in this study showed how such challenges can be handled to enable the cross-matching of single-cell epigenomic profiles. Notably, our method does not use distance-based measures (correlation or cosine distance) or hashing, latent-feature extraction approaches but leverages median expression and enrichment of top genes and peaks (MEXTEG and MESTEG) to mitigate batch and technical biases. Thus, the uniqueness of scEpiSearch lies in its large collection of reference cells and its statistical approach to reduce bias during the search for matching transcriptome and open-chromatin profiles. scEpiSearch enables single-cell search against a large pool of reference cell profiles, and provides low-dimensional embedding, summary word-cloud for overall notion about query scATAC-seq profiles and enrichment scores of genes to highlight possible markers for cell-types. Thus, scEpiSearch can also be useful for cross-validation of rare cellular states discovered using single-cell open chromatin profiles. The standalone version has an inbuilt processed reference and can be applied securely, locally, and confidentially towards sensitive and clinical data.

Our analysis revealed the benefits of mapping single-cell epigenomes to reference-cell profiles, as observed in the co-embedding of scATAC-seq and single-cell expression profiles, scEpiSearch based results outperformed integrative methods like Seurat, LIGER and Conos. Our analysis using scEpiSearch also highlighted a trend that using reference cell profiles for feature extraction and calculating distances among cells achieved better embedding of open-chromatin profiles than other latent feature extraction methods like SCALE, MINT and SCANORAMA. In addition, clustering based on similarity with reference cells could highlight new features in minor populations of cells that could have been overwhelmed by properties of cell-states in the majority due to the reference-free feature extraction method. While cells exist in a continuum of states across tumours and cell culture and actively respond to the environment, scEpiSearch was able to group these K562 and mESCs into discrete clusters based on single-cell accessibility peaks and similarity to reference datasets. We found a subset of K562 cells increasingly poised towards macrophages and dendritic lineage, indicative of regulatory re-wiring. We believe that the scEpiSearch comparison can help provide a better understanding of factors driving cellular heterogeneity. For example, we observed 4 mESC clusters based on chromatin accessibility after matching with reference datasets, where cluster-1 cells are similar to late blastocyst cells and likely with high cellular plasticity for response to stress. Some of the top enriched terms for mESC in cluster-1, like, unfolded protein response with stress in the endoplasmic reticulum (ER) (43), apoptosis, H4K5 and H4K8 acetylation and DNA damage, are known to co-occur(44, 45). Such results hint that heterogeneity in poising towards ER stress and unfolded protein response in multiple types of stem cells and preimplantation embryos (42) can also be studied using their scATAC-seq profile with scEpiSearch. There have also been multiple reports linking DNA damage, apoptosis, ER stress in the development of diabetes, cancer and other disorders(46, 47). Hence analysis of their single-cell open chromatin profiles using a pool of large reference cells with scEpiSearch could help better understand the cause or effect of such disorders.

Overall scEpiSearch can provide i) correct matching of query single-cell open chromatin profile to a large pool of single-cell profiles in a short time ii) cross-species search for query single-cell open-chromatin profile iii) correct co-embedding of single-cell open-chromatin profiles from two species iv) highlighting footprints within single-cells indicative of stress-response and apoptosis within a sub-population of embryonic stem cells. In future, we anticipate that scEpiSearch can help evaluate enhancer landscape activity within single-cells, incorporate increasingly single-cell histone modification datasets(48) (Supplementary Figure S8b) and provide an efficient search engine for major epigenetic regulators

## MATERIAL AND METHODS

### Preprocessing of single-cell ATAC-seq reference data

For each cell in the reference dataset of scATAC-seq, the read-count on every peak is normalized by its global accessibility score to enumerate its cell-type specificity. In other words, for every cell, peaks with cell-type-specific activity (possible enhancers) are highlighted by normalization with its global accessibility score. For both species, human and mouse, we have compiled global accessibility peak-list using several published open-chromatin profiles of bulk samples. For this purpose, we used the available peak lists of open chromatin profiles (DNAse-seq and ATAC-seq) of bulk samples, available at GEO database(49), UCSC genome browser(50) and iHEC portal(51). We merged the peaks lying within 1 kb of each other. The number of times a genomic site appeared as a peak in published open chromatin profiles of bulk samples was defined as its global accessibility score(52). Thus we have almost 1 million sites (width > 1 kbp) in our global accessibility list for both humans and mice. The normalized read-count *t_ij_* of a peak *i* in a single-cell *j* is calculated as

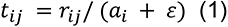

Here, the read-count *r_ij_* of the peak *i* in the single-cell *j* is normalized by its global accessibility score *a_i_* added with a pseudo-count *ε*.

Considering proximal (nearest) genes to all peaks as background, and genes nearest to top ten thousand enriched peaks are taken in the foreground set for every cell to calculate the p-value (of gene-enrichment) using Fisher’s exact test (based on hypergeometric distribution). Thus the equation of calculation of P-value of enrichment of genes can be written as :

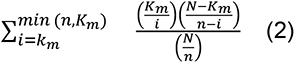

Here K_m_ represents the number of times a gene m appears in the background set, and N is the number of the appearance of all genes in the background. Whereas k_m_ is the number of times, the gene m appears in the foreground. At the same time, n is the total number of all genes in the foreground. Thus the genes proximal to many cell type specific sites (mostly enhancers) have a lower (more significant) p-value. Notice that it is the gene-enrichment calculation, and it is not the same as gene-set enrichment performed by different tools(39, 52). Using our approach we further processed large collection of single-cell open chromatin profile to make a in-built reference in scEpiSearch. For detail about processing of inbuilt reference epigenome profiles in scEpiSearch see supplementary Methods.

### Preprocessing reference single-cell expression

Datasets from different studies for human cells are assembled together using the same set of genes. Similarly, expression profiles of single-cells from mouse samples were assembled together. The single-cell RNA-seq based expression (FPKM) profiles were quantile normalized(53). Then for every gene, mean expression across all cells in reference is calculated. For every reference cell, the expression of a gene is normalized by its mean expression across all the cells. Thus, we convert absolute expression to cell-specific expression of genes for every cell. More detail about processing of reference single-cell RNA-seq data is mentioned in supplementary Methods

The framework of scEpiSearch is flexible enough to include very large reference single profiles. The current version of scEpiSearch consists of a compilation of single-cell expression profiles for 1288653 human cells and 2141797 mouse cells (total ∼3.3 million cells). Details about sources of single-cell RNA-seq profile is provided in supplementary table-1. The reference datasets of scATAC-seq profiles consist of 742297 human cells and 81173 mouse cells (total ∼800000 epigenome profiles). The information about sources of humans and mouse scATAC-seq profiles are provided in supplementary table-1.

### Query pre-processing

ScEpiSearch first highlights cell type specific peaks (mostly enhancers) by dividing the scATAC-seq read-count of every peak in the query peak-list by its global accessibility score (as shown in equation (1)). It finds genes proximal to peaks in the query scATAC-seq profile. In order to find proximal genes quickly, it first uses the pre-existing table of genes proximal to the peaks in the global accessibility list. For every query scATAC-seq profile, scEpiSearch finds overlap between its peaks and sites in the global accessibility list. With our analysis, we found that most of the time, scEpiSearch achieves overlap with the global accessibility list for 65-80% of the peaks in query. For peaks having an overlap with sites in the global accessibility list, ScEpisearch adapts their proximal genes from the pre-existing tables. For peaks, which do not have overlap with the global accessibility list, it searches proximal genes separately (if the accurate mode is selected). Thus, scEpiSearch avoids repeating the process of finding proximal genes for most of the peaks.

#### Gene enrichment scores of query

For every cell in the query, scEpiSearch selects genes proximal to peaks with high normalized read-count as foreground while keeping genes near all the peaks of query cell as background. For every query cell, it uses foreground and background genes to calculate gene-enrichment scores using Fischer’s exact test as explained in equation (2).

### Finding a match in scRNA-seq reference dataset

The approach of finding a matching expression profile to query scATAC-seq profile is based on the known fact that genes proximal to enhancers have high cell-type-specific expression. Therefore scEpiSearch first calculates gene enrichment scores of the query cells, which highlights genes frequently found near potential enhancers or cell-type specific sites. ScEpiSearch computes MExTEG (Median expression of top enriched genes) for query cells in representative expression vectors for clusters of RNA-seq cells, such that for query cell q the MExTEG value in reference cell m is

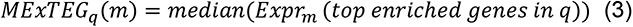

However, it is to be noticed that we do not use the raw expression (FPKM or TPM) of cells or representative of cells; rather, we use the cell-type-specific expression. In a cell, the cell-type-specific expression of a gene is calculated by dividing its expression with the median expression value of the same gene across all the cells. Thus for a reference scRNA-seq profile, MExTEG represents the median value of cell-type-specific expression of top enriched genes for query scATAC-seq profile. Since the search of matching expression profiles is done hierarchically, first, the MExTEG value is calculated using an expression vector for the cluster of reference scRNA-seq profiles. Notice that for a cluster, the representative expression vector contains the mean of cell-specific expression values of genes from cells belonging to that cluster. Further, the MExTEG for query cells is converted to P-value using the null model of representative expression vectors. Then top N clusters are chosen, which have the lowest p-value for MExTEG. Further, MExTEG for query cells is again computed using a cell-type-specific expression profile of single-cells present in top N selected clusters. One hundred cells are selected based on a higher MExTEG value. For 100 reference cells with high MExTEG for query, the MExTEG is converted to a p-value score using a null model. The p-value of these matches is calculated based on the null model explained below. As a final refinement score, a new P-value is calculated based on rank calculated using P-value-ranks. The details of the calculation of two types of P-values are provided below

#### Statistical approach to calculate significance of match

*A null model was prepared by* randomly selecting a few normalized scATAC-seq profiles using global accessibility scores of peaks. For the randomly selected 500 cells, the top thousand genes with the highest gene-enrichment score were extracted. Random pairs were made from selected 500 cells, and the top enriched gene lists of two cells in a pair were merged. Among the merged list of 2000 genes for every pair of cells, 1000 genes were randomly selected. Thus, we made thousand query vectors that served as false queries (cells), each having a thousand genes. For every cell in the set of false queries, MExTEG is calculated using cell-specific expression profiles of the reference cell to get a matrix of size 1000 x No of reference single-cells. For a reference cell, we calculate the p-value of similarity as the fraction of null model cells (false queries), which have higher values of MExTEG than for the query cell. Thus the P-value of match between query q and reference expression profile of cell m is calculated as

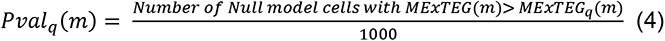

#### Rank based statistical approach to refine significance of match

In order to reduce bias in the search for matching cells, scEpiSearch further refines or adjusts the rank of matches due to P-values calculated using MExTEG. For this purpose, we keep the precalculated rank of every reference cell for all cells in the null model (false queries explained above). Such rank calculation provides a view of bias in the data and enumerates the number of times a reference cell comes in the top hit for the cells in the null model. Thus, after calculating the P-value of the match and determining the rank of a reference cell for a query cell, we calculate a new P-value. The new P-value of the match between a reference cell and a query cell is calculated as the fraction of cells in the null model for which the same reference cell has a better rank than for the query cell.

### Finding match among a huge set of reference single-cell epigenome profiles

Similar to the expression matching procedure, MESTEG (Median gene enrichment scores of top enriched genes) for query cells is calculated using representative GE (Gene enrichment) vectors for clusters of cells.

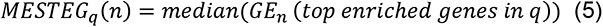

Further, the conversion of MESTEG value for the query to P-value is done using the null model for representative GE (Gene enrichment) vectors for clusters. After finding the top matching clusters using P-value for MESTEG, the matching is done at the single-cell level. Hence in the second round, MESTEG is calculated for query cells using all cells in top matching clusters. Again, these MESTEG values for the query are converted to p-values to find the significance of match with each reference cell in top matching clusters. Using the rank of matched reference cells, new p-value is calculated to reduce bias in search.

#### Statistical approach to calculate significance of match

Initially, a random selection of 500 normalized ATAC-seq cells GE scores are made from various profiles. Then, random pairs of cells were taken, and for every pair of cells, the mean normalized read-count was calculated. Thus, a total of one thousand false queries is made whose MESTEG is calculated for all scATACseq reference profiles. For every reference cell, the P-value of the match with the query is calculated as the fraction of null model cells (false queries), which have higher values of MESTEG than the query cell. Thus Pval for match between query q and reference cell n

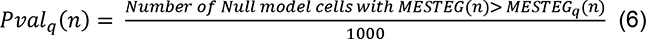

After converting MESTEG to p-value of the match, the rank of a reference cell for a query cell is used to calculate the new p-value. The new P-value of the match between a reference cell and a query cell is calculated as the fraction of cells in the null model for which the same reference cell has a better rank than for the query cell.

### Embedding of multiple Query scATAC-seq profile

We devised a novel method included in scEpiSearch to handle input multiple query read-count matrices with different peak-list from both human and mouse cells and perform their co-embedding. For this purpose, scEpiSearch computes gene enrichment scores for cells in each batch of query scATAC-seq profiles with different peaks/species separately. Since enrichment scores are calculated for all genes in same list, different scATAC-seq read-counts get transformed to the same feature set. Hence scEpiSearch integrates GE scores for different read-count matrices to the same matrix. However, scEpiSearch does not use GE directly for embedding as it is influenced by the batch effect. Once GE scores from multiple read-count matrices are assembled together, matches for queries are found in mouse single-cell expression profiles using the procedure described above. For mouse query cells, the null model for the mouse is used. For human query cells, scEpiSearch uses a null model made using human cells, just as explained above for cross-species search. After finding matching mouse cells for all query cells, scEpiSearch builds a network with every node representing a query cell. It connects two query cells (nodes) with an edge with a weight equal to the number of top matching reference cells belonging to the same cluster. For example, consider a query cell A for which 4 out of the top 10 matching mouse expression profiles belong to a sub-cluster X. If another query cell B has 5 out of the top 10 matching expression profiles from the same sub-cluster X., Then the weight of an edge between query cells A and B would be 4.

After calculating all edge weights, connections of a node with only K neighbours are kept.

Thus after making KNN based graph with edge weights, it uses Fruchterman & Reingold algorithm (see supplementary method) to calculate 2D coordinate of nodes. Fruchterman and Reingold algorithm(54) is a force-directed method of graph drawing (see Supplementary Methods). For utilization of the Fruchterman and Reingold algorithm to avoid overlap between dissimilar cells, we choose a proper value of K (number of neighbours) while making KNN based graph. After determining the 2D coordinate of nodes (representing query cells), it uses the networkX library (draw_network_nodes function) to plot the network for visualization (55). In order to assist users, it also performs spectral clustering (56) using the KNN based network of queries to find clusters of cells.

## EXPERIMENTAL METHODS

### Mouse embryonic stem cell culture

E14 mESC were cultured in 6-well dishes, pre-coated with 0.1% gelatin, and in Serum+LIF media containing DMEM knockout (Gibco #10829), 15% FBS (Gibco #10270), 1xPen-Strep-Glutamine (Gibco #10378), 1xMEM (Gibco #11140), 1xB-ME (Gibco #21985) and 1000 U/ml LIF (Merck #ESG1107).

## Acknowledgement and funding

We are thankful to the Indraprastha Institute of Information Technology - Delhi (IIIT-D) for the support of computational facilities and fellowship of PhD students. The research in KNN lab is supported by Villum Young Investigator grant (VYI#00025397), Novo Nordisk grants (#NNF18OC0052874) and Sino-Danish Centre (SDC).

## AUTHOR CONTRIBUTIONS

SM implemented the code and tested it and contributed to the writing of the manuscript. SC, NP and DS helped in the curation of the reference dataset and NP tested the public tools. KNN performed mESC experiments and scATAC-seq profiling used in this study. VK and KNN wrote the manuscript. VK also conceptualised the project.

## References

1. Buenrostro, J.D., Wu, B., Litzenburger, U.M., Ruff, D., Gonzales, M.L., Snyder, M.P., Chang, H.Y. and Greenleaf, W.J. (2015) Single-cell chromatin accessibility reveals principles of regulatory variation. Nature, 523, 486–490.

2. Corces, M.R., Shcherbina, A., Kundu, S., Gloudemans, M.J., Frésard, L., Granja, J.M., Louie, B.H., Eulalio, T., Shams, S. and Bagdatli, S.T. (2020) Single-cell epigenomic analyses implicate candidate causal variants at inherited risk loci for Alzheimer’s and Parkinson’s diseases. Nature genetics, 52, 1158–1168.

3. Lahnemann, D., Koster, J., Szczurek, E., McCarthy, D.J., Hicks, S.C., Robinson, M.D., Vallejos, C.A., Campbell, K.R., Beerenwinkel, N., Mahfouz, A. et al. (2020) Eleven grand challenges in single-cell data science. Genome Biol, 21, 31.

4. Lee, J.T.H., Patikas, N., Kiselev, V.Y. and Hemberg, M. (2021) Fast searches of large collections of single-cell data using scfind. Nature Methods, 18, 262–271.

5. Wu, K.E., Yost, K.E., Chang, H.Y. and Zou, J. (2021) BABEL enables cross-modality translation between multiomic profiles at single-cell resolution. Proceedings of the National Academy of Sciences, 118.

6. Wang, C., Sun, D., Huang, X., Wan, C., Li, Z., Han, Y., Qin, Q., Fan, J., Qiu, X. and Xie, Y. (2020) Integrative analyses of single-cell transcriptome and regulome using MAESTRO. Genome biology, 21, 1–28.

7. Jin, S., Zhang, L. and Nie, Q. (2020) scAI: an unsupervised approach for the integrative analysis of parallel single-cell transcriptomic and epigenomic profiles. Genome biology, 21, 1–19.

8. Danese, A., Richter, M.L., Chaichoompu, K., Fischer, D.S., Theis, F.J. and Colome-Tatche, M. (2021) EpiScanpy: integrated single-cell epigenomic analysis. Nat Commun, 12, 5228.

9. Stuart, T., Butler, A., Hoffman, P., Hafemeister, C., Papalexi, E., Mauck, W.M., 3rd, Hao, Y., Stoeckius, M., Smibert, P. and Satija, R. (2019) Comprehensive Integration of Single-Cell Data. Cell, 177, 1888–1902 e1821.

10. Liu, J., Gao, C., Sodicoff, J., Kozareva, V., Macosko, E.Z. and Welch, J.D. (2020) Jointly defining cell types from multiple single-cell datasets using LIGER. Nature protocols, 15, 3632–3662.

11. Barkas, N., Petukhov, V., Nikolaeva, D., Lozinsky, Y., Demharter, S., Khodosevich, K. and Kharchenko, P.V. (2019) Joint analysis of heterogeneous single-cell RNA-seq dataset collections. Nat Methods, 16, 695–698.

12. Luecken, M.D., Buttner, M., Chaichoompu, K., Danese, A., Interlandi, M., Mueller, M.F., Strobl, D.C., Zappia, L., Dugas, M., Colome-Tatche, M. et al. (2022) Benchmarking atlas-level data integration in single-cell genomics. Nat Methods, 19, 41–50.

13. Jin, W., Tang, Q., Wan, M., Cui, K., Zhang, Y., Ren, G., Ni, B., Sklar, J., Przytycka, T.M., Childs, R. et al. (2015) Genome-wide detection of DNase I hypersensitive sites in single cells and FFPE tissue samples. Nature, 528, 142–146.

14. Lai, B., Gao, W., Cui, K., Xie, W., Tang, Q., Jin, W., Hu, G., Ni, B. and Zhao, K. (2018) Principles of nucleosome organization revealed by single-cell micrococcal nuclease sequencing. Nature, 562, 281–285.

15. Cusanovich, D.A., Hill, A.J., Aghamirzaie, D., Daza, R.M., Pliner, H.A., Berletch, J.B., Filippova, G.N., Huang, X., Christiansen, L., DeWitt, W.S. et al. (2018) A Single-Cell Atlas of In Vivo Mammalian Chromatin Accessibility. Cell, 174, 1309–1324 e1318.

16. Fu, S., Wang, Q., Moore, J.E., Purcaro, M.J., Pratt, H.E., Fan, K., Gu, C., Jiang, C., Zhu, R. and Kundaje, A. (2018) Differential analysis of chromatin accessibility and histone modifications for predicting mouse developmental enhancers. Nucleic acids research, 46, 11184–11201.

17. Consortium, E.P. (2012) An integrated encyclopedia of DNA elements in the human genome. Nature, 489, 57–74.

18. Bujold, D., Morais, D.A.L., Gauthier, C., Cote, C., Caron, M., Kwan, T., Chen, K.C., Laperle, J., Markovits, A.N., Pastinen, T. et al. (2016) The International Human Epigenome Consortium Data Portal. Cell Syst, 3, 496–499 e492.

19. Han, X., Wang, R., Zhou, Y., Fei, L., Sun, H., Lai, S., Saadatpour, A., Zhou, Z., Chen, H., Ye, F. et al. (2018) Mapping the Mouse Cell Atlas by Microwell-Seq. Cell, 173, 1307.

20. Tetteroo, P.A., Massaro, F., Mulder, A., Schreuder-van Gelder, R. and von dem Borne, A.E. (1984) Megakaryoblastic differentiation of proerythroblastic K562 cell-line cells. Leuk Res, 8, 197–206.

21. Kleiveland, C.R. (2015) In Verhoeckx, K., Cotter, P., Lopez-Exposito, I., Kleiveland, C., Lea, T., Mackie, A., Requena, T., Swiatecka, D. and Wichers, H. (eds.), The Impact of Food Bioactives on Health: in vitro and ex vivo models, Cham (CH), pp. 161–167.

22. Jacoby, E., Nguyen, S.M., Fountaine, T.J., Welp, K., Gryder, B., Qin, H., Yang, Y., Chien, C.D., Seif, A.E. and Lei, H. (2016) CD19 CAR immune pressure induces B-precursor acute lymphoblastic leukaemia lineage switch exposing inherent leukaemic plasticity. Nature communications, 7, 1–10.

23. Slany, R.K. (2009) The molecular biology of mixed lineage leukemia. Haematologica, 94, 984.

24. Gallagher, R., Collins, S., Trujillo, J., McCredie, K., Ahearn, M., Tsai, S., Metzgar, R., Aulakh, G., Ting, R., Ruscetti, F. et al. (1979) Characterization of the continuous, differentiating myeloid cell line (HL-60) from a patient with acute promyelocytic leukemia. Blood, 54, 713–733.

25. Imaizumi, M., Uozumi, J. and Breitman, T.R. (1987) Retinoic acid-induced monocytic differentiation of HL60/MRI, a cell line derived from a transplantable HL60 tumor. Cancer research, 47, 1434–1440.

26. Kuleshov, M.V., Jones, M.R., Rouillard, A.D., Fernandez, N.F., Duan, Q., Wang, Z., Koplev, S., Jenkins, S.L., Jagodnik, K.M., Lachmann, A. et al. (2016) Enrichr: a comprehensive gene set enrichment analysis web server 2016 update. Nucleic Acids Res, 44, W90–97.

27. Sutherland, J.A., Turner, A.R., Mannoni, P., McGann, L.E. and Turc, J.M. (1986) Differentiation of K562 leukemia cells along erythroid, macrophage, and megakaryocyte lineages. J Biol Response Mod, 5, 250–262.

28. Sarna, M.K., Ingley, E., Busfield, S.J., Cull, V.S., Lepere, W., McCarthy, D.J., Wright, M.J., Palmer, G.A., Chappell, D., Sayer, M.S. et al. (2003) Differential regulation of SOCS genes in normal and transformed erythroid cells. Oncogene, 22, 3221–3230.

29. Fugazza, C., Barbarani, G., Elangovan, S., Marini, M.G., Giolitto, S., Font-Monclus, I., Marongiu, M.F., Manunza, L., Strouboulis, J. and Cantù, C. (2021) The Coup-TFII orphan nuclear receptor is an activator of the γ-globin gene. haematologica, 106, 474.

30. Zhao, C., Wang, B., Meng, D., Cao, Y., Yang, J., Zhao, X. and Chen, B. (2005) Study on rapid generation of dendritic cells from K562 cell line induced by A23187 alone. Zhonghua xue ye xue za zhi= Zhonghua Xueyexue Zazhi, 26, 408–412.

31. Hie, B., Bryson, B. and Berger, B. (2019) Efficient integration of heterogeneous single-cell transcriptomes using Scanorama. Nat Biotechnol, 37, 685–691.

32. Rohart, F., Eslami, A., Matigian, N., Bougeard, S. and Le Cao, K.A. (2017) MINT: a multivariate integrative method to identify reproducible molecular signatures across independent experiments and platforms. BMC Bioinformatics, 18, 128.

33. Lopez, R., Regier, J., Cole, M.B., Jordan, M.I. and Yosef, N. (2018) Deep generative modeling for single-cell transcriptomics. Nat Methods, 15, 1053–1058.

34. Xiong, L., Xu, K., Tian, K., Shao, Y., Tang, L., Gao, G., Zhang, M., Jiang, T. and Zhang, Q.C. (2019) SCALE method for single-cell ATAC-seq analysis via latent feature extraction. Nat Commun, 10, 4576.

35. Ester, M., Kriegel, H.-P., Sander, J. and Xu, X. (1996), kdd, Vol. 96, pp. 226–231.

36. Granja, J.M., Klemm, S., McGinnis, L.M., Kathiria, A.S., Mezger, A., Corces, M.R., Parks, B., Gars, E., Liedtke, M., Zheng, G.X.Y. et al. (2019) Single-cell multiomic analysis identifies regulatory programs in mixed-phenotype acute leukemia. Nat Biotechnol, 37, 1458–1465.

37. Buenrostro, J.D., Corces, M.R., Lareau, C.A., Wu, B., Schep, A.N., Aryee, M.J., Majeti, R., Chang, H.Y. and Greenleaf, W.J. (2018) Integrated single-cell analysis maps the continuous regulatory landscape of human hematopoietic differentiation. Cell, 173, 1535–1548. e1516.

38. Pliner, H.A., Packer, J.S., McFaline-Figueroa, J.L., Cusanovich, D.A., Daza, R.M., Aghamirzaie, D., Srivatsan, S., Qiu, X., Jackson, D., Minkina, A. et al. (2018) Cicero Predicts cis-Regulatory DNA Interactions from Single-Cell Chromatin Accessibility Data. Mol Cell, 71, 858–871 e858.

39. McLean, C.Y., Bristor, D., Hiller, M., Clarke, S.L., Schaar, B.T., Lowe, C.B., Wenger, A.M. and Bejerano, G. (2010) GREAT improves functional interpretation of cis-regulatory regions. Nature biotechnology, 28, 495–501.

40. Lastun, V.L., Grieve, A.G. and Freeman, M. (2016), Seminars in cell & developmental biology. Elsevier, Vol. 60, pp. 10–18.

41. Bellazzo, A., Di Minin, G. and Collavin, L. (2017) Block one, unleash a hundred. Mechanisms of DAB2IP inactivation in cancer. Cell Death & Differentiation, 24, 15–25.

42. Yang, Y., Cheung, H.H., Tu, J., Miu, K.K. and Chan, W.Y. (2016) New insights into the unfolded protein response in stem cells. Oncotarget, 7, 54010.

43. Lin, T., Lee, J.E., Kang, J.W., Shin, H.Y., Lee, J.B. and Jin, D.I. (2019) Endoplasmic reticulum (ER) stress and unfolded protein response (UPR) in mammalian oocyte maturation and preimplantation embryo development. International journal of molecular sciences, 20, 409.

44. Bolland, H., Ma, T.S., Ramlee, S., Ramadan, K. and Hammond, E.M. (2021) Links between the unfolded protein response and the DNA damage response in hypoxia: a systematic review. Biochemical Society Transactions.

45. Dhar, S., Gursoy-Yuzugullu, O., Parasuram, R. and Price, B.D. (2017) The tale of a tail: histone H4 acetylation and the repair of DNA breaks. Philosophical Transactions of the Royal Society B: Biological Sciences, 372, 20160284.

46. Yoshida, H. (2007) ER stress and diseases. The FEBS journal, 274, 630–658.

47. Urra, H., Dufey, E., Avril, T., Chevet, E. and Hetz, C. (2016) Endoplasmic reticulum stress and the hallmarks of cancer. Trends in cancer, 2, 252–262.

48. Rotem, A., Ram, O., Shoresh, N., Sperling, R.A., Goren, A., Weitz, D.A. and Bernstein, B.E. (2015) Single-cell ChIP-seq reveals cell subpopulations defined by chromatin state. Nature biotechnology, 33, 1165–1172.

49. Barrett, T., Suzek, T.O., Troup, D.B., Wilhite, S.E., Ngau, W.-C., Ledoux, P., Rudnev, D., Lash, A.E., Fujibuchi, W. and Edgar, R. (2005) NCBI GEO: mining millions of expression profiles—database and tools. Nucleic acids research, 33, D562–D566.

50. Lee, C.M., Barber, G.P., Casper, J., Clawson, H., Diekhans, M., Gonzalez, J.N., Hinrichs, A.S., Lee, B.T., Nassar, L.R. and Powell, C.C. (2020) UCSC Genome Browser enters 20th year. Nucleic acids research, 48, D756–D761.

51. Bujold, D., de Lima Morais, D.A., Gauthier, C., Côté, C., Caron, M., Kwan, T., Chen, K.C., Laperle, J., Markovits, A.N. and Pastinen, T. (2016) The international human epigenome consortium data portal. Cell systems, 3, 496–499. e492.

52. Chawla, S., Samydurai, S., Kong, S.L., Wu, Z., Wang, Z., Tam, W.L., Sengupta, D. and Kumar, V. (2021) UniPath: a uniform approach for pathway and gene-set based analysis of heterogeneity in single-cell epigenome and transcriptome profiles. Nucleic acids research, 49, e13–e13.

53. Cole, M.B., Risso, D., Wagner, A., DeTomaso, D., Ngai, J., Purdom, E., Dudoit, S. and Yosef, N. (2019) Performance Assessment and Selection of Normalization Procedures for Single-Cell RNA-Seq. Cell Syst, 8, 315–328 e318.

54. Fruchterman, T.M. and Reingold, E.M. (1991) Graph drawing by force-directed placement. Software: Practice and experience, 21, 1129–1164.

55. Schult, D.A. and Swart, P. (2008), *Proceedings of the 7th Python in science conferences (SciPy 2008)*. Pasadena, CA, Vol. 2008, pp. 11–16.

56. Xiang, T. and Gong, S. (2008) Spectral clustering with eigenvector selection. Pattern Recognition, 41, 1012–1029.

57. Chen, X., Miragaia, R.J., Natarajan, K.N. and Teichmann, S.A. (2018) A rapid and robust method for single cell chromatin accessibility profiling. Nature communications, 9, 1–9.

58. Xu, W., Wen, Y., Liang, Y., Xu, Q., Wang, X., Jin, W. and Chen, X. (2021) A plate-based single-cell ATAC-seq workflow for fast and robust profiling of chromatin accessibility. Nature protocols, 16, 4084–4107.

